# Computational and Experimental Identification of Potential Neutralizing Peptides Derived from Human ACE2 Against SARS-CoV-2 Infection

**DOI:** 10.1101/2025.09.01.673483

**Authors:** Qiaobin Yao, Vidhyanand Mahase, Wangheng Hou, Ruth Cruz-Cosme, Qiyi Tang, Shaolei Teng

## Abstract

The human angiotensin-converting enzyme 2 (hACE2) is the primary receptor for the entry of SARS-CoV-2. Some human alleles of ACE2 exhibit an improved affinity for the SARS-CoV-2 Spike protein. However, the impact of ACE2 polymorphisms on SARS-CoV-2 infection remains unclear. Our previous study predicted that G431 and S514 in the receptor-binding domain (RBD) od SARS-CoV-2 S1 domain are important for S protein stability, and that S protein residues G496 and F497 and ACE2 residues D355 and Y41 are critical for the RBD–ACE2 interaction (1). In this study, we explored the potential of hACE2-derived neutralizing peptides as a therapeutic strategy against SARS-CoV-2 and investigated how ACE2 polymorphisms affect RBD–ACE2 binding affinity. We applied computational saturation mutagenesis to systematically screen the binding affinity changes among all possible ACE2 missense mutations within the ACE2–Wuhan-S1 complex. Mutations at ACE2 residues D355 and Y41 were predicted to weaken binding affinity, whereas those at N330 and D30 enhanced it. We identified six ACE2 regions (19–49, 65–102, 320–333, 348–359, 378–395, 552–563) to be vital for ACE2–RBD interaction. We synthesized peptides corresponding to these six regions and tested them using a pseudotyped viral particle system and dot blot assay. Three peptides were confirmed to bind with S protein, and four exhibited inhibitory effects. We aligned ACE2–Wuhan-S1 and ACE2–Omicron-S1 complexes, conducted correlation analysis, and observed similar binding patterns, suggesting that these peptides also have potential to neutralize Omicron strains.

**IMPORTANCE:** SARS-CoV-2 continues its global spread. In this research, we identified six regions within ACE2 that are vital for interaction with the viral S RBD and have potential to neutralize SARS-CoV-2 infection. Among the six peptides derived from ACE2, three were confirmed to bind with S protein of Wuahan strain, and four exhibited inhibitory effects on Wuahn strain SARS-CoV-2. We also found ACE2 residues D355 and Y41 as weakening affinity, and N330 and D30 as enhancing it. We also aligned this complex with the ACE2–Omicron-S1 complex, performed correlation analyses, and compared their patterns of stability changes upon mutations, and obtained similar results, indicating that these peptides may also be effective against Omicron variants. These results provide insight into the role of ACE2 polymorphism in viral entry and suggest that hACE2-derived peptides may offer a promising therapeutic strategy against SARS-CoV-2, demonstrating strong consistency between our computational predictions and experimental outcomes.

## INTRODUCTION

The COVID-19 pandemic, caused by the severe acute respiratory syndrome coronavirus 2 (SARS-CoV-2), has led to unprecedented global health and economic crises (2-5). The World Health Organization (WHO) reported more than 778 million confirmed cases and nearly 7 million deaths due to SARS-CoV-2 infection as of July 2025 (6). SARS-CoV-2 is the seventh identified human coronavirus. Coronaviruses (CoVs) are a group of large and enveloped RNA viruses carrying a single-stranded positive-sense RNA genome (7). The S glycoprotein facilitates the initial stages of viral entry, including attachment to the host cell membrane and membrane fusion. The host proteases cleave the S protein into two subunits, S1 and S2. The S1 subunit is accountable for receptor Angiotensin-converting enzyme 2 (ACE2) recognition, while the S2 subunit is responsible for membrane fusion (8, 9). Spike-ACE2 interaction is required for the infection of SARS-CoV-2, so we hypothesize that blocking the interaction will effectively prevent and treat SARS-CoV-2-caused disease.

There exist two major types of peptides that have been reported to be effective in preventing SARS-CoV-2 infection: ACE2-derived (10) and spike-derived (11). Peptide inhibitors derived from ACE2 with high affinity for SARS-CoV-2 could potentially address the challenges associated with antibody-based therapies, including issues in manufacturing, distribution, and drug delivery. In fact, ACE2 serves as a common receptor for several coronaviruses, positioning it as an attractive target for developing broad-spectrum coronavirus therapies that can resist viral mutations and enhance preparedness for future pandemics (12). And we hypothesize that ACE2-derived peptides not only have anti-SARS-CoV-2 activity but also present low or no toxicity.

Previous studies have primarily focused on characterizing the SARS-CoV-2 Spike-ACE2 interaction through molecular simulations and structural analyses or developing ACE2-mimicking peptides as therapeutic candidates. While these approaches have identified key binding residues, they often lack a systematic strategy for optimizing peptide sequences to improve therapeutic efficacy (13, 14). Given the lack of systematic optimization in prior approaches, computational tools can offer a rational approach to peptide design by identifying optimal modification sites to enhance stability and reduce degradation (15, 16).

In this study, we combined computational and experimental approaches to develop anti-SARS-CoV-2 peptides. Six ACE2-derived peptides were designed and tested with a pseudotyped viral system, demonstrating that these peptides effectively reduced viral entry in a cell culture system. This approach enables a more precise and efficient peptide design strategy, overcoming key limitations in existing ACE2-based therapeutics and offering a promising framework for the development of stable and effective antiviral peptides.

## RESULTS

### Identification of ACE2 regions that bind to SARS-CoV-2 spike protein

To generate required information supporting the development of ACE2-derived neutralizing peptides against SARS-CoV-2, we computationally evaluated the effects of 113,324 non-redundant missense mutations in the ACE2 protein within the ACE2-Wuhan-S1 complex (PDB: 6LZG). Specifically, we calculated the binding affinity changes (ΔΔΔG) resulting from these mutations. A heatmap of ΔΔΔG values is shown in Figure 1A, while Figure 1B presents the mean values of ΔΔΔG at each ACE2 residue position, along with ΔΔΔG value of substitutions to alanine.

**Fig. 1.**
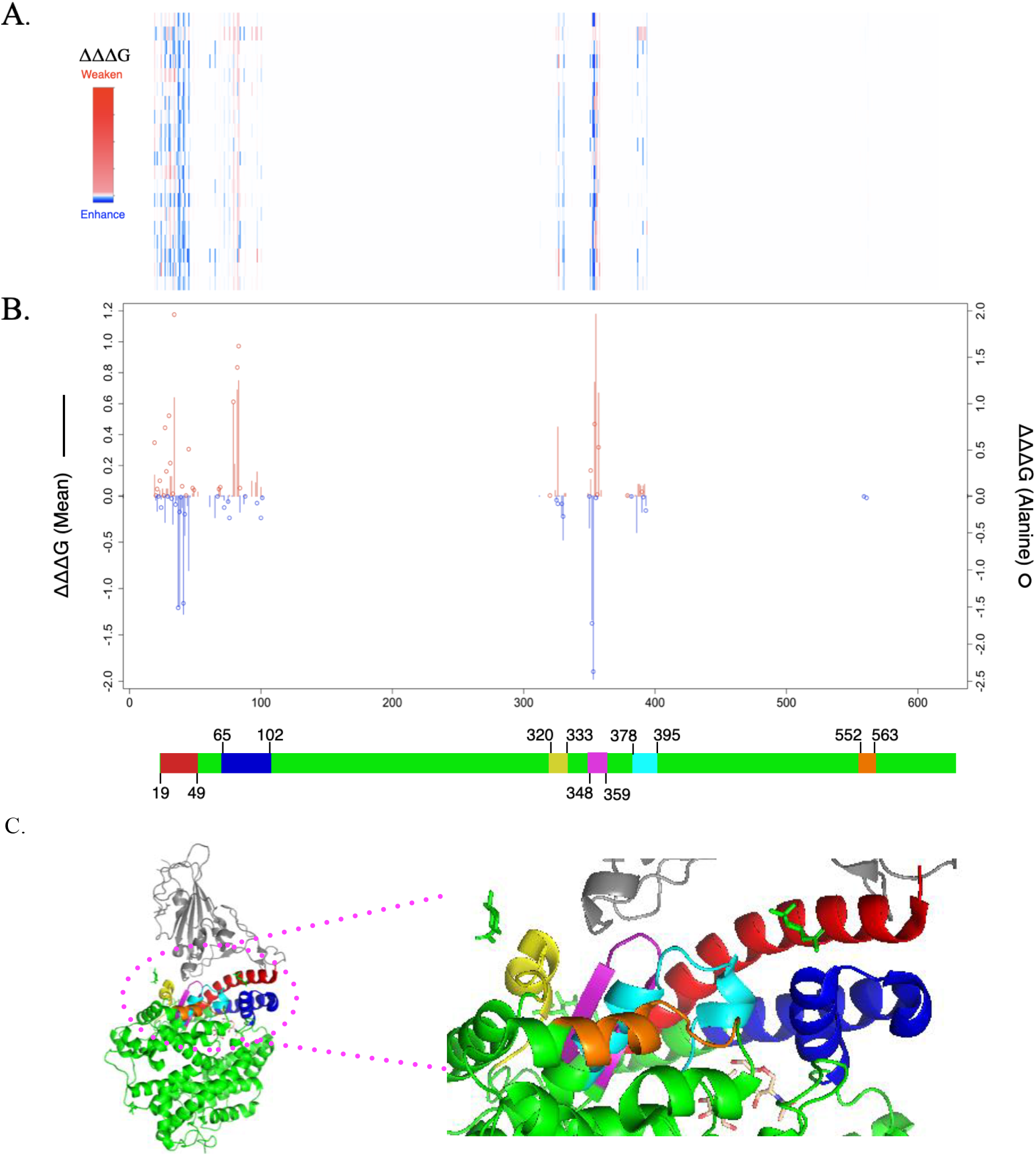
Effects of residues and mutations on ACE2-Wuhan-S1 interaction. (1) The line chart summarizes the binding energy changes for ΔΔΔG mean of residues (2) and ΔΔΔG of substitutions to Alanine (circle) in ACE2 residues, a decrease in binding affinity (red) and an increase in binding affinity (blue). (1) Heatmaps show the ΔΔΔG of all ACE2 mutations. (C) Six peptide regions are shown in PyMol images. ACE2 is shown in green. Wuhan-Hu-S-1 is shown in gray. Region 1(19-49): Red; Region 2 (65-102): Blue; Region 3(320-333): Yellow; Region 4 (348-359): Purple; Region 5 (378-395): Cyan; Region 6 (552-563): Orange.

Our analysis identified six key ACE2 regions-residues 19-49, 65-102, 320-333, 348-359, 378-395, 552-563-that are associated with significant changes in binding affinity. These regions appear to play a critical role in stabilizing the interaction between the human ACE2 and the SARS-CoV-2 S1 protein. Given their importance in receptor-virus binding, these ACE2 regions represent promising candidates for the design of ACE2-derived peptides capable of competitively inhibiting SARS-CoV-2 by mimicking native ACE2 and blocking the spike protein’s receptor-binding domain. Such peptides may offer an effective therapeutic strategy against SARS-CoV-2 infection.

### Effects of ACE2 mutations on ACE2-Wuhan-S1 Interaction

To determine how ACE2 mutations affect the ACE2-Wuhan-S1 interaction, we selected key ACE2 residues whose alterations significantly influence binding affinity. Based on the mean values of ΔΔΔG, five residues-R357, M82, G354, Y83, and D355-were found to enhance binding affinity, while five other residues-K353, G352, Y41, E37, and D38-reduced the binding affinity. Among these, residue K353 showed the strongest enhancing effect (ΔΔΔG mean = -1.89 kcal/mol), with K353Y mutation exhibiting greatest reduction in binding free energy (ΔΔΔG =-4.3 kcal/mol), indicating a significant increase in ACE2-Wuhan-S1 binding. Conversely, residue D355 had the strongest weakening effect (ΔΔΔG mean = 1.18 kcal/mol), with D355W mutation yielding the highest ΔΔΔG value at 5.86 kcal/mol, reflecting weakened interaction (Fig. 2A, Table 1).

**Table 1.** Effects of key mutations on ACE2–SARS-Cov-2 S interaction.

| ACE2-Wuhan-S1 complex |  |  | ACE2-Omicron-S1 complex |  |  |
| --- | --- | --- | --- | --- | --- |
| Variant | $\Delta\Delta\Delta G$ | rs ID | Variant | $\Delta\Delta\Delta G$ | rs ID |
| R357Y | 2.25 | \ | D355F | 5.09 | \ |
| M82A | 1.39 | \ | Y41G | 2.94 | \ |
| G354T | 3.28 | \ | Y83G | 1.88 | \ |
| Y83W | 2.23 | \ | G354P | 4.88 | \ |
| D355W | 5.86 | \ | E35K | 3.35 | rs1348114695 |
| K353Y | -4.3 | \ | H34W | -3.39 | \ |
| G352L | -3.02 | \ | L39E | -2.2 | \ |
| Y41H | -2.57 | rs768024356 | F72W | -1.84 | \ |
| E37R | -2.47 | \ | K31L | -2.17 | \ |
| D38Q | -2.41 | \ | D30L | -2043 | \ |
| K26R | 0.01 | rs4646116 | K26R | -0.02 | rs4646116 |
| K341R | 0 | rs138390800 | K341R | 0.07 | rs138390800 |
| S19P | 0.88 | rs73635825 | S19P | 0.4 | rs73635825 |
| E37K | -0.83 | rs146676783 | E37K | 0.33 | rs146676783 |
| M82I | 0.42 | rs766996587 | M82I | 0.36 | rs766996587 |

**Fig. 2.**
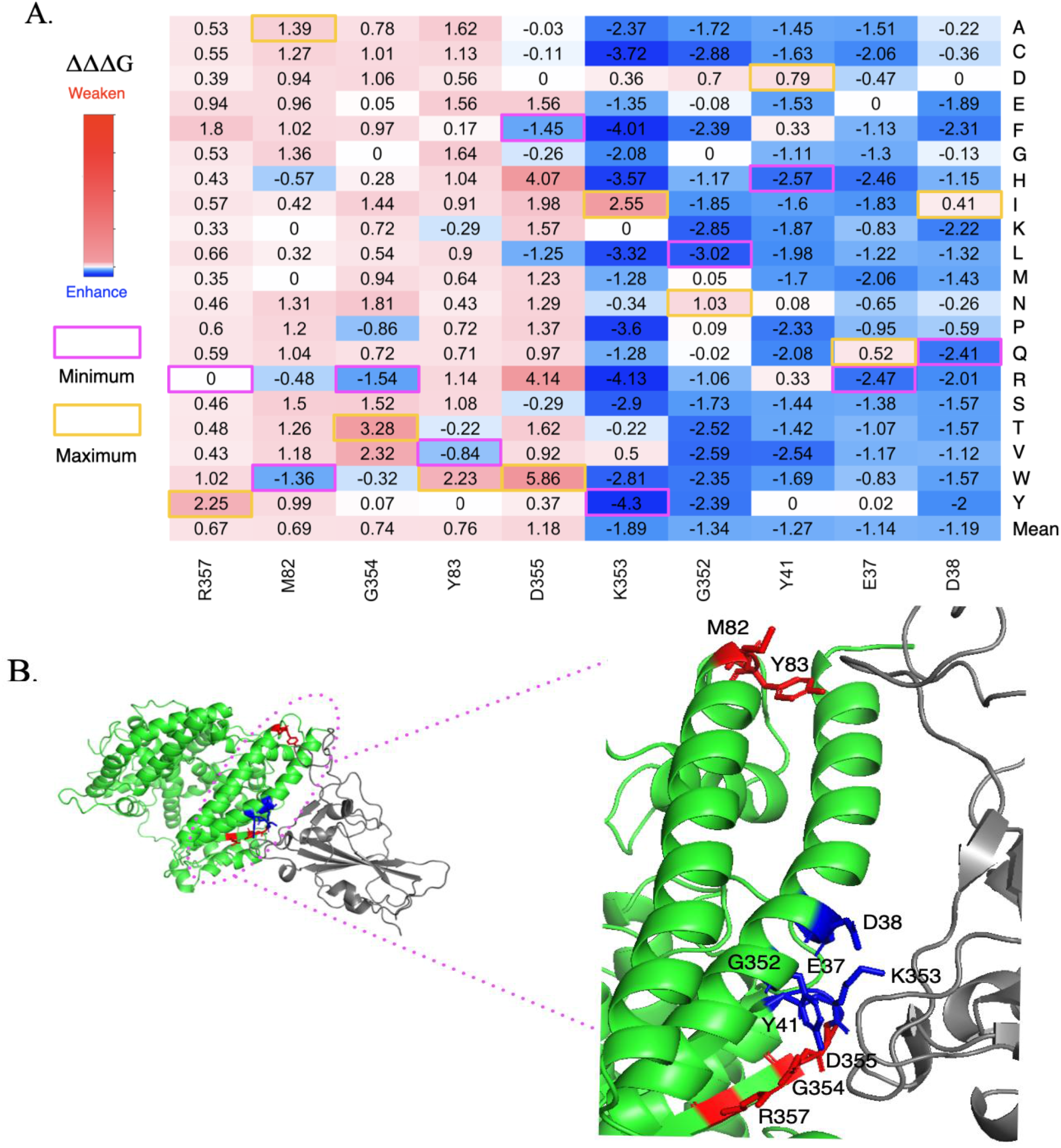
Key residues that affect ACE2 – Wuhan-S1 binding affinity. (1) heatmaps are shown as maximum values (yellow), and minimum (magenta); (1) Key residues are shown in blue or red in PyMol images. ACE2 is shown in green. Wuhan-S1 is shown in gray.

As shown in Fig. 2B, all those residues are located at the binding interfaces of the ACE2-Wuhan-S1 complex and participate in critical interaction network. Notably, residues R357, G354, D355, K353, Y41, E37, and D38 have been previously reported as key residues in both ACE2–SARS-CoV-2 S RBD and ACE2–SARS-CoV S RBD interfaces (1, 17). D30, K31, and K353 interact with the K417, E484, and L452 of SARS-CoV-2 S protein, and the mutations on those residues may facilitate antibody escape. Y41 interacts with the N501Y of SARS-CoV-2 S protein, and both the ACE2 Y41 mutations of and the S N501Y mutation have been shown to enhance ACE2-Wuhan-S1 binding affinity (18, 19).

Moreover, we found that the residues with significant effects on binding affinity are clustered within these specific regions (Fig. 1 and 2). E37, D38, and Y41 are located in the 1^st^ region (19-49); M82 and Y83 in the 2^nd^ region (65-102); and G352, K353, G354, D355, and R357 in 4^th^ region. In addition, we investigated the binding energy change induced by human genetic variations. The common variants, E37K and G352V, can enhance the ACE2-Wuhan-S1 interaction. In contrast, variants M82I and D355N can weaken the ACE2-Wuhan-S1 interaction.

### Experimental validation of hACE2-derived peptides binding to SARS-CoV-2 Spike protein

To confirm the binding affinity of the six hACE2-derived peptides to the Spike protein of SARS-CoV-2, we conducted an *in vitro* binding assay followed by a dot blot analysis. First, each peptide dissolved in RPMI-1640 was spotted onto an NC membrane at equal amounts, after naturally drying, the membrane was visualized to capture the red spots as shown in Fig. 3A, showing an equal amount of peptides spotted onto the membrane. The membrane was then fixed and subsequently incubated with a S protein-containing solution. Following three washes with PBS-T, the membrane was probed with HRP-labeled anti-Spike antibody. Detection of bound S protein was performed by dot blot assay. The results showed strong binding of the S protein to peptides ACE19-49 and ACE65-102, and weak affinity with ACE348-359 (Fig. 3B). The remaining peptides showed no detectable binding. Quantitative analysis revealed that the relative binding affinities of ACE19-49 and ACE65-102 were significantly higher than that of a random control peptide (Fig. 3C). These findings demonstrate that the ACE2-S binding predicted by the computational modeling requires experimental confirmation.

**Fig. 3.**
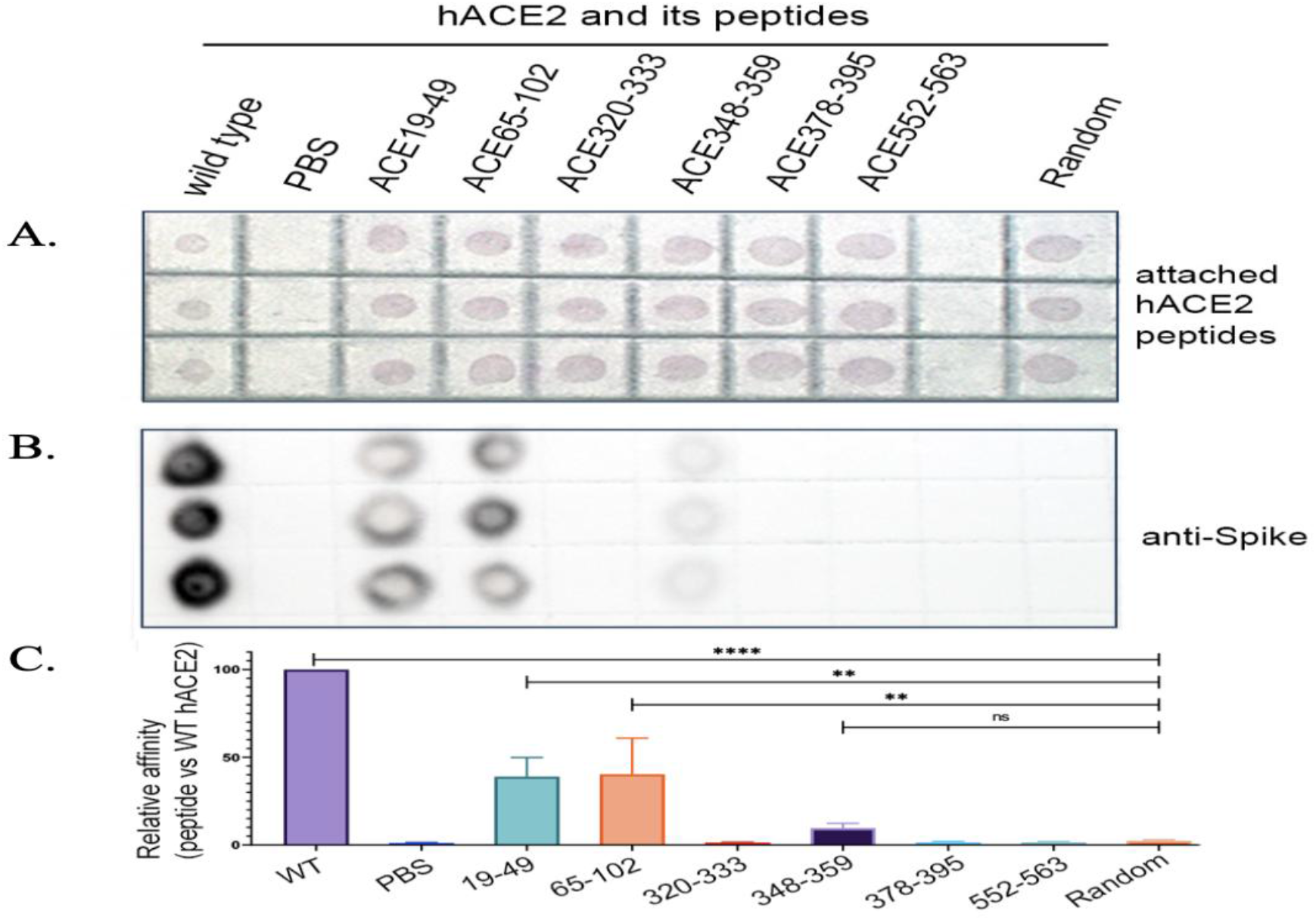
*In vitro* binding assay followed by a dot blot analysis. (1) Each peptide was spotted onto an NC membrane (red); (1) Detection of bound S protein was performed by dot blot assay; (C) Quantitative analysis.

### ACE2-derived peptides significantly inhibit SARS-CoV-2 infection

To evaluate whether hACE2-derived peptides inhibit SARS-CoV-2 infection, we employed a pseudotyped viral particle system to circumvent the need for a BSL-3 facility required for live SARS-CoV-2. This system allowed us to assess the inhibitory effects of the peptides in a cell culture model. As shown in Fig. 4, a 2-fold serial dilution of each peptide mixed with the pseudotyped SARS-CoV-2 particle and used to infect BHK-hACE2 cells. All six hACE2-derived peptides significantly reduced Pseudotyped SARS-CoV-2 infection in a dose-dependent manner. Notably, the two peptides with the strongest Spike protei-binding affinities exhibited the most potent inhibitory effects, with IC50 values of 6.59 nM and 2.23 nM, respectively. Interestingly, the peptide of ACE378-395, despite its low binding affinity, also demonstrated strong inhibition of viral infection. This discrepancy suggests that additional mechanisms may contribute to its inhibitory effect and warrants further investigation. These findings demonstrate that the computationally predicted hACE2-derived peptides can effectively attenuate SARS-CoV-2 infection in hACE2-expressing cells. Moreover, our established pseudotyped virus system provides a valuable platform for studying the inhibitory effects of potential therapeutic agents against SARS-CoV-2.

**Fig. 4.**
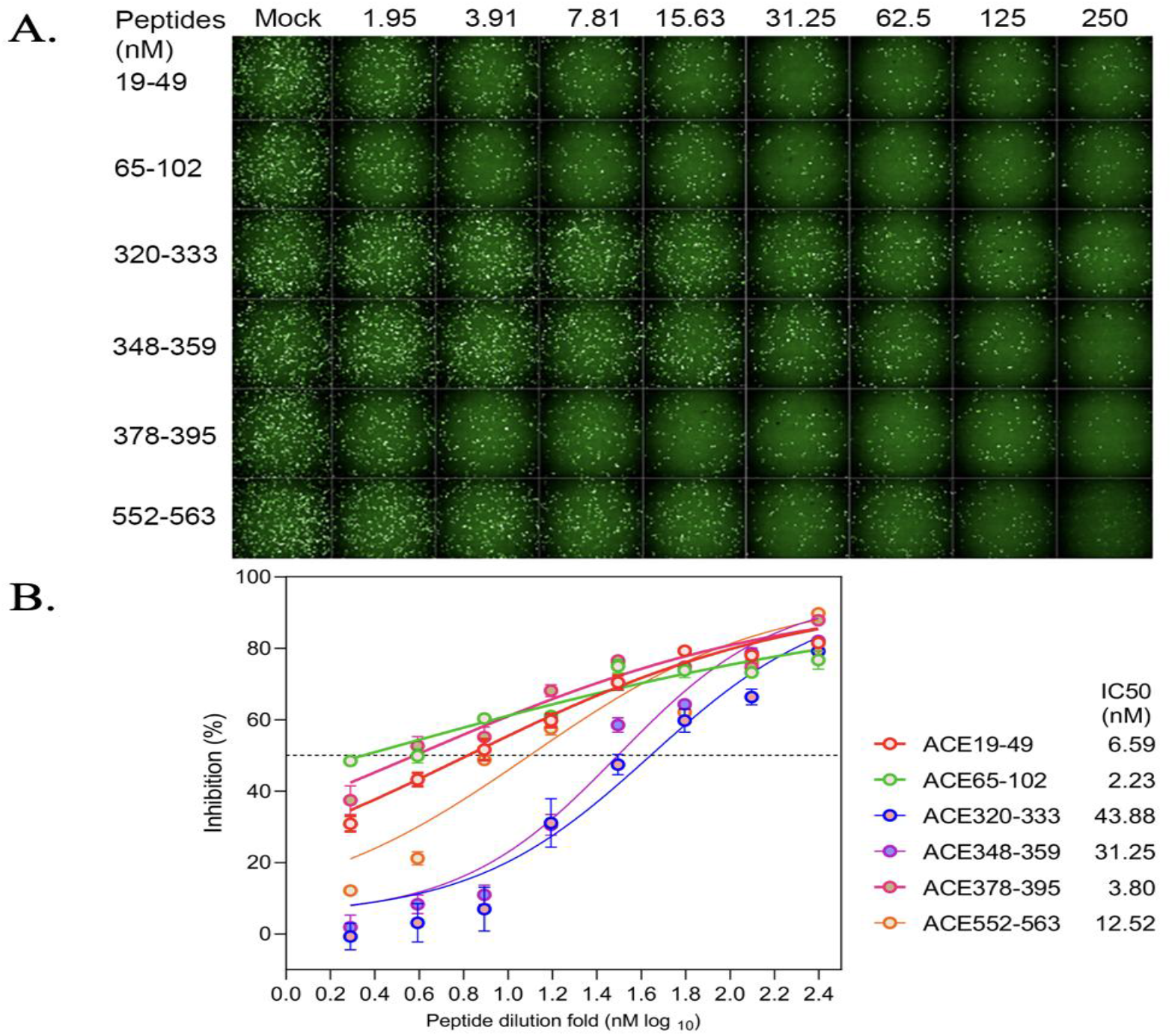
The inhibitory effects of the peptides: using pseudotyped viral particle system to evaluate the inhibitory effects of the peptides in a cell culture model.

### Comparison of ACE2-Wuhan-S1 and ACE2-Omicron-S1 complexes

To investigate whether these potential ACE2-neutralizing peptides can also be applicable to the ACE2-Omicron-S1 complex, we performed structural alignment of the ACE2 chains of both the ACE2-Wuhan-S1 and ACE2-Omicron-S1 complexes (Fig. 5A) and compared ΔΔΔG values of ACE2 mutations in both complexes (Fig. 5B, 5C). Comparing the root mean square deviation (RMSD) of corresponding atoms in two proteins is a standard measure to evaluate their similarity. Lower RMSD values indicate greater similarity. The RMSD value between the two complexes is 637, which indicates a high degree of structural similarity between ACE2 conformations in both complexes (Fig. 5A). We then compared the ΔΔΔG values of ACE2 mutations in the ACE2-Wuhan-S1 and ACE2-Omicron-S1 complexes (Fig. 5B), and observed a modest correlation in the binding affinity effects across both complexes (*R* = 0.23). Structural alignment and correlation analysis revealed consistent patterns in how ACE2 mutation affect binding in both ACE2-Wuhan-S1 and ACE2-Omicron-S1 complexes. The mean values of ΔΔΔG at each ACE2 residue position in both complexes are shown in the line charts of Fig. 5C. As shown in the combined line chart, although some top residues differed, five ACE2 regions with potential as neutralizing peptides overlapped almost completely between the two complexes.

**Fig. 5.**
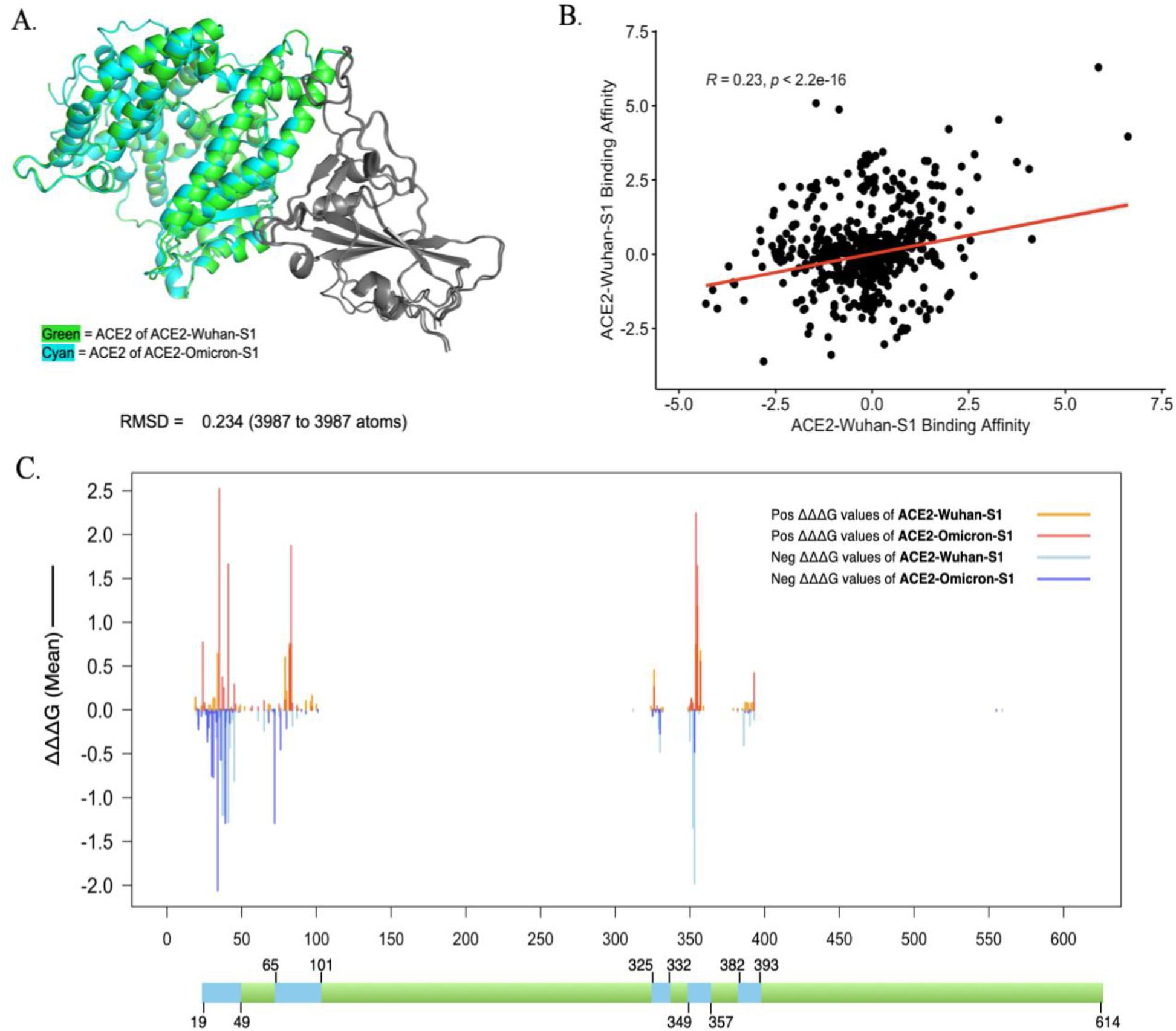
Comparison of ACE2-Wuhan-S1 and ACE2-Omicron-S1 complexes. (1) Structural alignment of ACE2-Wuhan-S1 and ACE2-Omicron-S1 complexes. (1) Correlation analysis of ACE2-Omicron-S1 and ACE2-Wuhan-S1 binding affinity ΔΔΔG. (C) Combined line chart of the mean binding affinity ΔΔΔG.

## DISCUSSION

SARS-CoV-2 infection is initiated by the binding of the viral spike protein to the host receptor ACE2, a critical step that governs viral entry and pathogenesis (8). Genetic variation in ACE2 has been reported to alter binding affinity with the spike receptor-binding domain (RBD) (20-22), yet its implications for viral susceptibility and therapeutic intervention remain incompletely understood. ACE2-derived peptides offer the potential for broad-spectrum inhibition of coronaviruses (13, 14) while minimizing issues associated with antibody-based therapies (15, 16). Our study systematically investigated ACE2 polymorphisms and designed ACE2-derived peptides to evaluate their capacity to interfere with SARS-CoV-2 entry, which bridged computational predictions with experimental validation.

This study focuses on investigating the potential of ACE2-based peptides and analyzed key residues in ACE2 proteins that are important for its interaction with the S protein, based on the binding affinity changes caused by ACE2 missense mutations. We used the ACE2-Wuhan-S1 complex as a reference to identify potential peptide candidates. First, a computational saturation mutagenesis aimed at discovering ACE2-derived peptides capable of neutralizing SARS-CoV-2 identified five regions within the ACE2 protein (residues 19–49, 65–101, 325–332, 349–357, and 382–393) that are highly likely to interact with the S protein. Next, the affinities of these peptides to the spike protein were tested *in vitro*, and three out of five showed higher affinities. Additionally, we used a pseudotyped viral particle system to test the peptides’ effects on the infection of SARS-CoV-2, and the result showed that the ACE2-derived peptides significantly attenuated the viral infection in hACE2-complemented cells, suggesting the potential of computationally identified peptides in reducing the severity of SARS-CoV-2 infection. Furthermore, we performed a structural alignment between the ACE2-Wuhan-S1 and ACE2-Omicron-S1 complexes to assess variations in their binding interfaces. To quantitatively compare the binding affinity between these two complexes, we conducted a correlation analysis, followed by a comparative examination of line charts illustrating the binding trends. Our results demonstrated a high degree of similarity between the two complexes, suggesting that the identified peptides may also have the potential to neutralize SARS-CoV-2 Omicron variants.

We analyzed the human ACE2 genetic variants, as these polymorphisms can influence ACE2 structure, stability, expression, and its binding affinity to the SARS-CoV-2 spike protein, thereby affecting individual susceptibility and disease severity. The prevalence of ACE2 genetic variants differs across populations, potentially contributing to disparities in COVID-19 outcomes (23, 24). Notably, variants such as rs4646116 (K26R) and rs138390800 (K341R) are associated with reduced protein stability and are more common in certain populations (25). Several known ACE2 variants, including rs73635825 (S19P), rs146676783 (E37K), and rs766996587 (M82I), showed distinct effects across viral strains (Table 1). For instance, the M82I and S19P variants decreased binding affinity in the Omicron complex but had minimal impact in the Wuhan complex. Conversely, the E37K variant increased binding in the Wuhan interaction (ΔΔΔG = –1.83 kcal/mol) but reduced it in Omicron (ΔΔΔG = 0.22 kcal/mol), emphasizing the variant-specific nature of ACE2–spike interactions. We further examined the impact of specific ACE2 mutations on spike binding. Within the ACE2–Wuhan-S1 complex, mutations at D355, Y83, and G354 significantly weakened binding affinity, while K353, G352, and Y41 enhanced it (Fig. 2A). Interestingly, the Y41 mutation exhibited opposite effects between variants, enhancing ACE2– Wuhan-S1 binding (ΔΔΔG = –1.27 kcal/mol) but weakening binding in the ACE2–Omicron-S1 complex (ΔΔΔG = 1.88 kcal/mol).

This study investigates the development of targeted therapeutic strategies by identifying key residues and mutations that affect the ACE2 protein’s binding with the SARS-CoV-2 S1 protein. In particular, we highlight neutralizing peptides that can block this critical interaction. Our computational analyses offer valuable insights into the systemic effects of ACE2 mutations on binding, supporting the rational design of effective treatments and vaccines to help mitigate the ongoing impact of COVID-19.

## MATERIALS AND METHODS

### Structure preparation

To investigate the impact of mutations on the interaction between Omicron-S and hACE2, we conducted a comparison between ACE2-Omicron-S1 (PDB ID: 7WBP), a crystal structure that illustrates the binding of Omicron-S to ACE2, and ACE2-Wuhan-S1 (PDB ID: 6LZG), which reveals the configuration of RBD and RBM proteins that facilitate viral entry. The structures were acquired from the Protein Data Bank (26).

### Mutation collection

To assess the impact of mutations on proteins, we applied the computational saturation mutagenesis to mutate all residues in the corresponding structures to all other 19 amino acid types. This approach entails introducing mutations at these specific locations to incorporate all conceivable amino acids. Subsequently, the resulting variants are examined to identify any potential alterations in protein stability or interactions with other proteins (27). Specifically, we employed an in-house Perl script to produce the list of all possible mutations in each residue. Then, we used the lists in the FoldX (28) program to simulate the effect of the mutation on the intricate protein structure. We collected ACE2 genetic variations in different populations from two data sources: gnomAD v4.3 (29) and HGMD (30).

### Energy calculations

Before doing any energy calculations, all protein structures were fixed using the ‘RepairPDB’ command on the command line interface in FoldX offcial website. This command operates by altering certain residues to reduce the overall free energy of the protein structure. The FoldX program was utilized for interaction analysis and computing saturation mutagenesis, total energy calculations, Van der Waals interactions, hydrogen bonding, and other energy correlations.

The calculation of binding energy change (ΔΔΔG) was performed using FoldX for each mutation. The ‘AnalyseComplex’ command was utilized to determine the free folding energy change, specifically the binding energy change between the contacts. We calculated the ΔΔG(binding) by breaking down each protein and analyzing their energies. To get the change in binding free energy (ΔΔΔG), we subtract the ΔΔG(binding) value from the total energy of the protein complex, specifically from the wild-type complexes. The mathematical equation provides the value of the change in the binding free energy (ΔΔΔG) between the mutant structure and the wild-type structure.

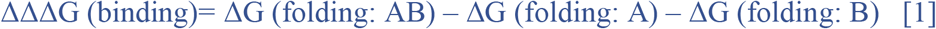

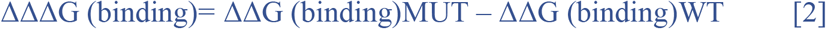

A negative value of ΔΔΔG indicates an increased binding affinity, whereas a positive value denotes a decrease in binding affinity. Using the obtained ΔΔΔG values, we employed the R program to create heatmaps and line graphs (31).

### Structural alignments and correlation analysis

We also used PyMoL (32) to do the Structural Alignments between ACE2-Wuhan-S1 (PDB ID: 6LZG) and ACE2-Omicron-S1 (PDB ID: 7WBP) by using the PyMol software’s ‘fetch and align’ commands, as well as display their key mutations, resulting in the generation of visual representations of 3-D structures. Also, we did the correlation analysiscompared to compare the dddG values of those two structures by using R package (31). Also, R was used to generate graphs and perform analysis of variance test and *t-*test in statistical comparisons of energy changes.

### Binding assays of Spike-hACE2 and Spike-hACE2-derived peptides

A dot blot assay was performed to evaluate the bindings of hACE2 and its derived peptides to SARS-CoV-2 Spike protein. Briefly, hACE2 and the peptides (synthesized and purified by Sino Biological and Sangon Biotech, Shanghai) were dissolved in RPMI-1640 medium (no serum) and spotted onto nitrocellulose membrane (13) at 0.5 μg per dot. After air drying, an initial photograph was taken to capture the red color of phenol red in RPMI-1640 (Fig. 3A). The membrane was fixed with 4% paraformaldehyde at room temperature for 10 min, washed with PBS-T (PBS containing 0.05% Tween-20), and air-dried at room temperature. Subsequently, the membrane was incubated with PBS containing 5% skim milk for 1 hour at room temperature with gentle shaking. After 3 washes with PBS-T, the membrane was incubated with the Spike protein solution (0.1mg/mL) for 1 hour at room temperature with gentle shaking. After 3 washes with PBS-T, the membrane was treated with an anti-Spike antibody conjugated with horseradish peroxidase (HRP) (2F9-HRP, New England Biolabs). Positive signals were detected using the ChemiDoc MP System (Bio-Rad) with the SuperSignal West Femto Maximum Sensitivity Substrate. The grayscale intensities of the dots were quantified using ImageJ software.

### Inhibitory effects of the hACE2-derived peptides on SARS-CoV-2 infection

A pseudotyped viral particle system was used to avoid using the actual viral particle that needs a BSL-3 facility. The Pseudotyped ΔG-GFP (G*ΔG-GFP) rVSVw/ pCAGGS-G-Kan system was purchased from Kerafast Company (https://www.kerafast.com/productgroup/171/pseudotyped-g-gfp-gg-gfp-rvsv). ΔG-GFP is a modified form of vesicular stomatitis virus (rVSV) that is limited in its ability to replicate and the infectivity of these pseudotyped viruses is limited to a single round of replication, they can be handled using standard BSL-2 containment practices.

The spike gene of the Wuhan-Hu-1 strain (GenBank: MN908947) was codon-optomized for expression in human cells and cloned into the eukaryotic expression plasmid pCAG to create pCAG-nCoV-S. The plasmid pCAG-nCoV-S was introduced into Vero-E6 cells and incubated for 48 hours. Subsequently, the VSVdG-EGFP-G virus (33) was inoculated to the cells expressing SARS-CoV-2 spike protein for 1 hour. Then, VSVdG-EGFP-G virus-containing supernatant was replaced with medium containing anti-VSV-G rat serum that blocks the infection of residual rVSVdG-G. The supernatant was collected at 24 hours post rVSVdG-G infection and centrifuged and filtered (0.45-μm pore size) to remove cell debris, aliquoted and stored at −80°C for use. The synthetic hACE2-derived peptides were diluted in a continuous gradient twofold from 250 nM to 1.95 nM and mixed with VSV-SARS-CoV-2-S virus (MOI=0.05) and incubated at 37°C for 1 hour. The mixtures were added to BHK21-hACE2 cell for 12 hours. Fluorescent assays were performed to visualize the infected cells (GFP).

## ACKNOWLEDGMENTS

This study was supported by the National Science Foundation (NSF) under award DBI-2000296 and partially by NSF award HRD-2406155, as well as the National Institute on Minority Health and Health Disparities of the National Institutes of Health (NIH) under award 2U54MD007597. Any opinions, findings, and conclusions or recommendations expressed in this study are those of the authors and do not necessarily reflect the views of the NSF or NIH.

